# Transformation and allelic exchange in *Orientia tsutsugamushi*

**DOI:** 10.64898/2026.06.22.733791

**Authors:** Paige E. Allen, Jason R. Hunt, Travis J. Chiarelli, Jason A. Carlyon

## Abstract

*Orientia tsutsugamushi* is a mite-transmitted obligate intracellular bacterium that causes the potentially deadly zoonosis, scrub typhus. The absence of genetic tools for *Orientia* have limited studies of the microbe-host interactions that underlie scrub typhus. To address this gap, we developed a protocol for transforming and achieving allelic exchange in *O. tsutsugamushi* str. Ikeda. From evaluating multiple cell lines and antibiotics, we found that contact-inhibited EA.hy926 human endothelial-like cells best supported *Orientia* replication and that chloramphenicol was an effective selection marker. We engineered a homologous recombination cassette encoding a codon-modified version of the *O. tsutsugamushi ank13* gene (OTT_RS04140) (CM*ank13*) and its promoter alongside genes for mScarlet-I and chloramphenicol acetyltransferase under control of the *O. tsutsugamushi tsa22*-up and *tsa56*-down promoters, respectively. A PCR product encompassing the cassette and chromosomal flanking regions was transformed into *O. tsutsugamushi* via electroporation or CaCl_2_, the latter of which better preserved bacterial and host cell viability. EA.hy926 cells inoculated with transformed *O. tsutsugamushi* were grown in glass-bottom plates in the presence of chloramphenicol and imaged by live-cell microscopy to identify cultures containing mScarlet-I positive bacteria, which could be maintained in perpetuity. Chromosomal integration of the CM*ank13* cassette and loss of wild-type *ank13* were verified by PCR and nanopore sequencing. This report establishes platforms for genetically manipulating *O. tsutsugamushi* and building additional genetic tools to investigate this globally significant pathogen.

**IMPORTANCE:** *Orientia tsutsugamushi* causes scrub typhus, a globally emerging rickettsiosis that can have a high mortality rate and has been a known human disease since the fourth century. Of the genera of obligate intracellular bacterial pathogens that cause human disease, *Orientia* is the only one for which genetic tools have not been developed. This has limited understanding of *O. tsutsugamushi*-host dynamics that drive the bacterium’s pathobiology and hindered development of novel treatment or protection strategies against scrub typhus. Here, we successfully transformed and achieved allelic exchange in *O. tsutsugamushi*. Transgenic bacteria were selected via antibiotic resistance, validated by PCR and nanopore sequencing, and visualized by immunofluorescence and live-cell fluorescence imaging. Our report includes detailed descriptions of empirically determined host cell cultivation, multiplicity of infection, transformation, and selection conditions to provide a foundation on which other researchers can build. Overall, this work begins to establish a genetic toolbox for *O. tsutsugamushi*.

## INTRODUCTION

*Orientia tsutsugamushi* is a mite-transmitted obligate intracellular bacterium that invades leukocytes and endothelial cells to cause scrub typhus, a life-threatening infection for which no vaccine exists. The disease afflicts over one million people annually in the Asia-Pacific (1). Non-travel related cases also occur in several African countries (2–6), Chile (7), Peru (8), and the United Arab Emirates (9). The detection of *Orientia* DNA in mites in North Carolina and seropositivity against *O. tsutsugamushi* antigens in individuals from this region indicates the presence of *Orientia* spp. in the United States (10–12). Scrub typhus has a 6% case fatality rate that can be much higher in the absence of timely treatment. Even with antibiotic therapy, the fatality rate remains approximately 1.4%. Like other rickettsioses, its initial presentation as an undifferentiated febrile illness can confound diagnosis and delay proper treatment allowing the infection to progress with the development of pneumonitis, acute respiratory distress, acute renal failure, meningoencephalitis, and myocarditis (1, 13). Heavy *O. tsutsugamushi* burdens in the cytosol of endothelial cells is linked to widespread vasculitis that can culminate in hypotensive shock, systemic vascular collapse, multiorgan failure, and death (1, 14).

The earliest written description of scrub typhus was in China in 313 AD, making it a known human disease for nearly two millennia (15). Despite numerous and varied attempts over the past 80-plus years, an effective scrub typhus vaccine has not been produced (13). In terms of better understanding *Orientia*-host interactions that drive pathogenesis, leveraging the availability of several annotated *O. tsutsugamushi* strain genomes and using them in RNAseq and comparative virulence analyses has proven informative (16–23). The use of surrogate systems, biochemical studies of recombinant *Orientia* proteins, and functionally interrogating ectopically expressed versions of *Orientia* proteins that recapitulate infection-associated phenomena in trans have also advanced understanding of *O. tsutsugamushi* pathobiology (18, 24–37). Still, the *O. tsutsugamushi* factors that facilitate invasion of host cells, intracellular replication, virulence, and pathogenesis, particularly those that could be targeted to treat or prevent scrub typhus, remain inadequately defined. These knowledge gaps are due, in part, to *O. tsutsugamushi* genetic intractability. Of the six obligate intracellular bacterial pathogen genera that afflict humans, *Orientia* is the only one for which genetic tools have not been developed (38, 39).

The *O. tsutsugamushi* chromosome is AT-rich (∼69%) and ranges from 1.9 to 2.3 Mb among strains, nearly twice that of other obligates (16, 21–23, 40). Forty to 50% of the genome consists of repetitive sequences and mobile genetic elements including more than 200 Tc-1/mariner transposons. The Himar1 transposase, a member of the Tc-1 mariner superfamily, has been extensively used to generate random transposon insertion mutants for many obligate intracellular bacterial species (16, 21–23, 40–54). However, due to the extensive presence of Tc-1/mariner transposable elements throughout the *O. tsutsugamushi* chromosome, introduction of Himar1 transposase would likely induce multiple unwanted insertion events per bacterium, making this approach impractical. As an alternative consideration, endonucleases, which act as a defense mechanism against foreign DNA, are not predicted to be carried by *O. tsutsugamushi* (16, 21–23, 40). Therefore, DNA delivered into the organism for mutagenesis may not be at risk for restriction endonuclease mediated damage.

Here, we report the first successful method for transformation and allelic exchange in *O. tsutsugamushi*. We determined the minimum bacteria-to-host cell ratio required to achieve a productive infection and empirically evaluated host cell options, cultivation and antibiotic selection conditions, and DNA delivery methods. Leveraging this information, we transformed *O. tsutsugamushi* with a DNA cassette encoding an *Orientia* codon-modified gene, fluorescent reporter, and selection marker each under the control of a distinct promoter. We verified cassette integration into the *O. tsutsugamushi* chromosome by PCR and sequencing and employed live-cell microscopy to identify host cells infected with fluorescent transgenic *O. tsutsugamushi* that were expanded. Transgenic *Orientia* could be maintained with or without antibiotic selection. Overall, this study establishes methods for genetically manipulating this neglected pathogen.

## RESULTS

### Selection of a host cell line and cultivation conditions that maximize *O. tsutsugamushi* growth

Experiments in this study were performed using the *O. tsutsugamushi* Ikeda strain, hereafter referred to as *O. tsutsugamushi*, a clinical isolate that causes severe disease in humans and is lethal in laboratory mice (17, 55–58). Because transformation of obligates tends to be inefficient and post-transformation recovery of transgenic organisms typically takes multiple weeks (38, 39), we needed to use an immortalized host cell line that would robustly support but not outpace *O. tsutsugamushi* intracellular replication. We evaluated three that are amenable to *O. tsutsugamushi* infection: EA.hy926 (human endothelial-like), HeLa (human cervical epithelial), and Vero (African green monkey kidney) cells (24, 25, 59, 60). EA.hy926 and Vero cells exhibit contact inhibition while HeLa cells do not. Each was maintained in medium supplemented with 1% or 10% fetal bovine serum (FBS) and synchronously infected at a multiplicity of infection (MOI) of 5–10. Over a 72-h period, which allowed for maximal replication prior to bacterial egress and host cell lysis (25), *O. tsutsugamushi* genomic equivalents (GE) were quantified by qPCR. Host cell proliferation was measured as the reduction in CellTrace Far Red signal using flow cytometry. The *Orientia* DNA load was greatest in HeLa cells grown in 10% and 1% FBS (Fig. 1A-C). However, EA.hy926 cells grown in 10% FBS exhibited the most restricted cell proliferation (Fig. 1D-G).

**FIG 1.**
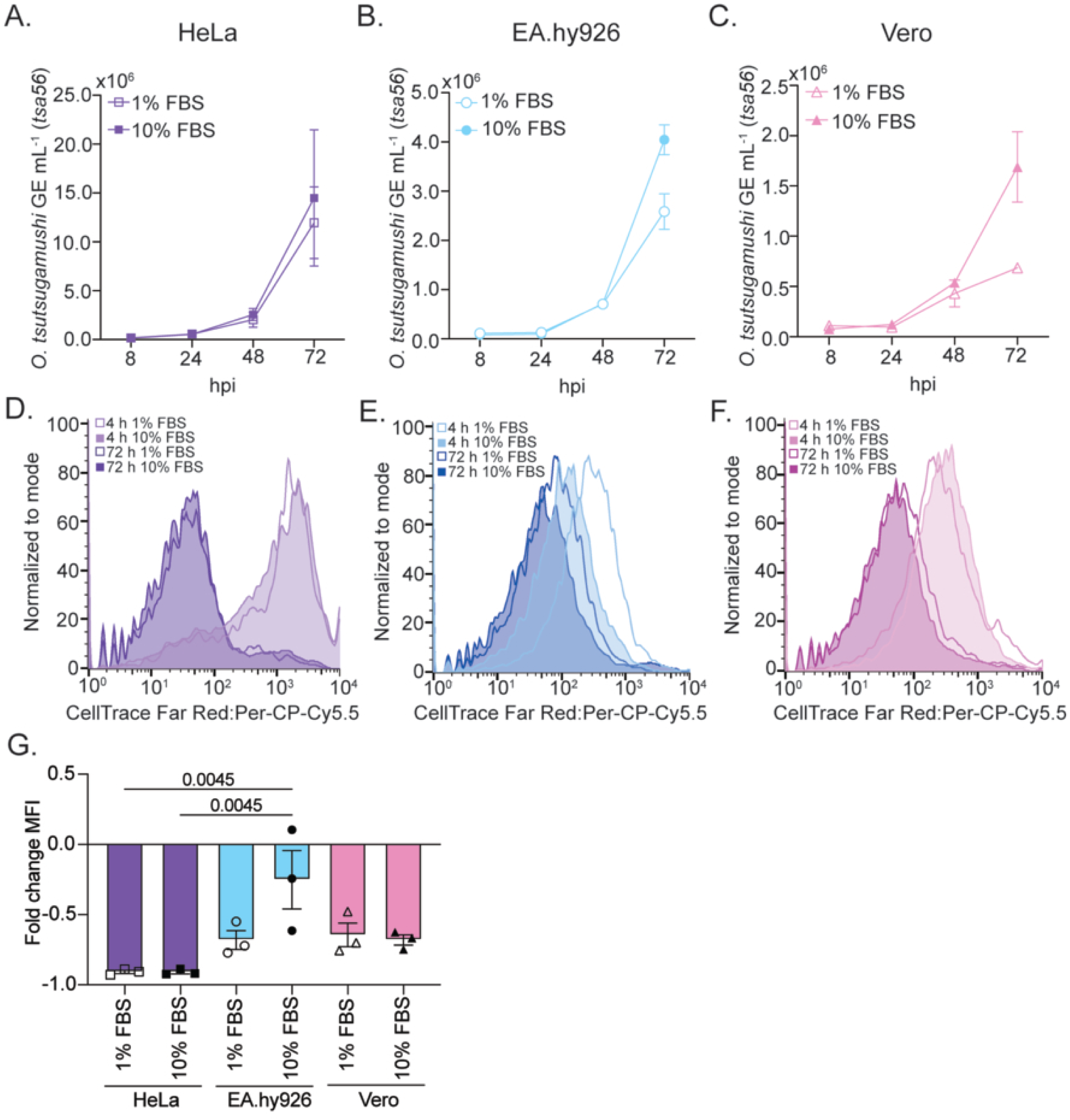
Determining the optimal cell line model for *O. tsutsugamushi* maintenance. HeLa (A, D), EA.hy926 (B, E), or Vero (C, F) cells were infected with *O. tsutsugamushi*, and the medium was supplemented with 1% or 10% FBS. (A–C) Total DNA isolated at 8, 24, 48, and 72 hpi was subjected to qPCR using primers targeting *O. tsutsugamushi tsa56* to determine the *Orientia* GE/mL. Data are means ± SD and representative of three experiments with similar results. (D–F) Cells were incubated with CellTrace Far Red following by examination of host cell proliferation flow cytometry. (G) The fold-change in mean fluorescent intensity (MFI) was determined by normalizing each sample to the four h timepoint. Data are presented as the means ± SEM and are representative of 3 experiments with similar results. One-way ANOVA with Tukey’s post hoc test was used to test for a significant difference among groups. Statistically significant values are indicated.

Isolation of isogenic mutants of obligate intracellular microbes requires successful expansion from a single clone, which is often achieved using a plaque assay (38, 39, 59, 60). We employed a plaque assay similar to that developed for *O. tsutsugamushi* infected murine embryonic cells by Hanson (61) using infected HeLa and EA.hy926 cells cultivated in media containing 10% or 1% FBS. Due to the robust growth of HeLa cells even when supplemented with 1% FBS, we included the eukaryotic protein synthesis inhibitor, cycloheximide as an additional condition. Plates were monitored daily. No plaques were observed by 14 d post-infection (dpi) for any condition regardless of whether plates were left unstained or stained with either of the non-lethal reagents, neutral red or 3-(4,5-dimethylthiazol-2-yl)-2,5-diphenyltetrazolium bromide (MTT). Overall, of the conditions tested, EA.hy926 cells cultured in 10% FBS most favorably support *in vitro O. tsutsugamushi* infection and will be used to recover transgenic bacteria post-transformation, but a plaque assay will not be employed further in this study.

### Determination of the minimum MOI that yields a productive *O. tsutsugamushi* infection

Although an MOI of 5–10 yielded productive *O. tsutsugamushi* infections, we rationalized that this MOI could be difficult to achieve with transformed bacteria given that the methods proven to be successful for other rickettsial organisms compromise viability. We considered it prudent to determine the minimum MOI that would lead to a productive infection. Host cell-free *O. tsutsugamushi* bacteria were recovered from heavily infected HeLa cells via mechanical disruption using zirconia beads. Confluent EA.hy926 monolayers paired in triplicate in individual wells of a 96-well plate were incubated with *O. tsutsugamushi* to achieve an MOI of 5–10 and five-fold serial dilutions thereof. The MOI in which both infected and uninfected cells were counted was verified by immunofluorescence microscopy at 4 h post-infection (pi). Total DNA isolated from inocula and infected cells at 72 hpi was subjected to qPCR to quantify *Orientia* GE. The undiluted inoculum yielded an MOI of 7 with 96.6% of the cells being infected at 4 h (Fig. 2). Consistent with previous reports, this translated to a 1.15-log increase in the bacterial load at 72 h. MOIs of 1.3 and 0.53 led to slightly lower yields. An MOI of 0.3 with only 26.7% of infected cells at 4 h led to no bacterial expansion. Given that an MOI of 0.53 with 46.7% of cells infected was the most diluted condition that yielded a productive infection, we infer that the required minimum *O. tsutsugamushi*-to-host cell ratio is greater than one. Moreover, undiluted inocula contained an average of 1.5 x 10^6^ *O. tsutsugamushi* GE mL^-1^ while inocula that failed to yield a productive infection had an average of 9 x 10^3^ *O. tsutsugamushi* GE mL^-1^. Thus, approximately 0.6% of the inocula generated using our isolation procedure is either not viable or incompetent for infection. Overall, these data indicate that our *O. tsutsugamushi* isolation procedure retains near complete bacterial viability and establish that an MOI > 1 will be needed post transformation to recover transgenic *O. tsutsugamushi* mutants.

**FIG 2.**
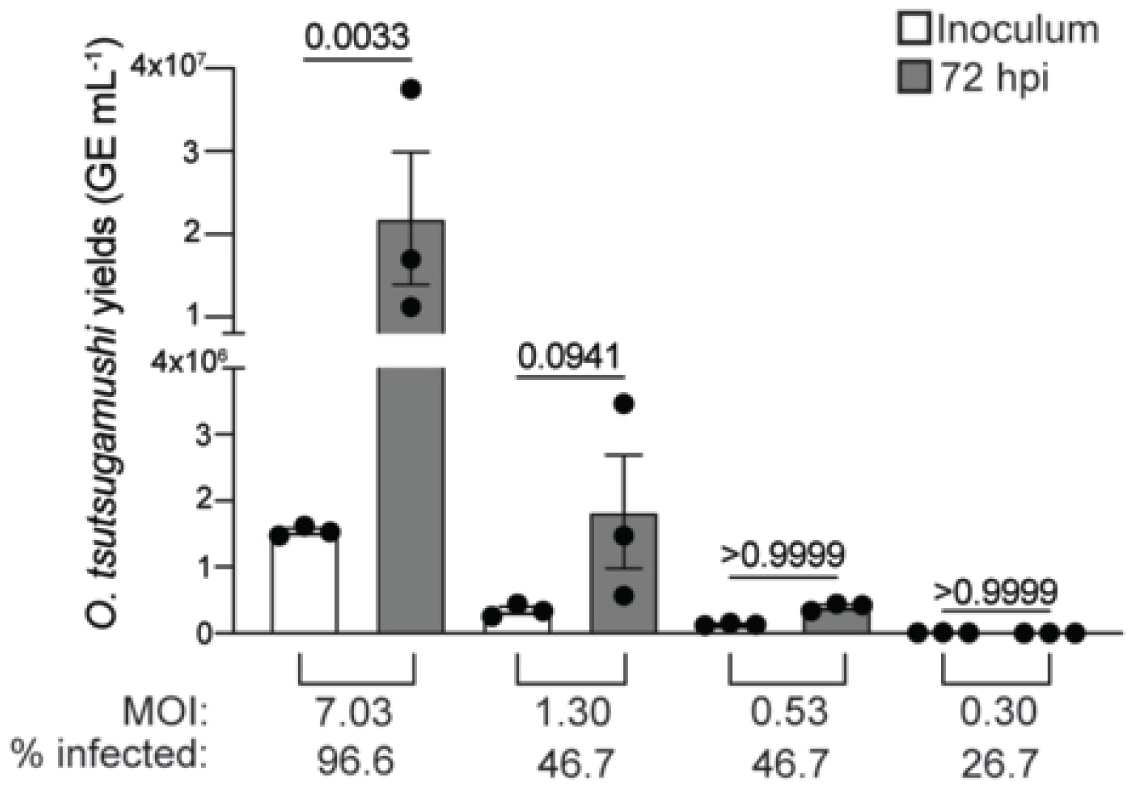
Defining the minimum MOI required for a productive *O. tsutsugamushi* infection. EA.hy926 cells were incubated in 96-well plates with *O. tsutsugamushi* to achieve an MOI of 5–10 and five-fold serial dilutions thereof. The mean number of bacteria per cell (MOI) and percentage of infected cells (% infected) were quantified by immunofluorescence microscopy. Total DNA was collected at the time of inoculation and at 72 h post infection and subjected to qPCR using primers targeting *O. tsutsugamushi tsa56* to determine the *Orientia* GE/mL. Data are and representative of three experiments with similar results. One-way ANOVA with Tukey’s post hoc test was used to test for a significant difference among groups. Statistically significant values are indicated.

### Identification of potential selection agents for *O. tsutsugamushi*

Selection of genetically manipulated bacteria typically relies on the introduction of antibiotic resistance markers during transformation. Chloramphenicol, rifampicin, spectinomycin, and gentamycin have each been successfully employed as selection agents for obligate intracellular bacteria (38). Based on genome analysis, *O. tsutsugamushi* is predicted to be naturally resistant to gentamicin and spectinomycin (62), although this has yet to be experimentally examined. *O. tsutsugamushi* clinical isolates are susceptible to chloramphenicol and rifampicin (63). However, effective concentrations of either drug for halting *O. tsutsugamushi* replication while maintaining host cell viability had not been determined for EA.hy926 cells. Synchronously infected EA.hy926 cells were treated with each of the four antibiotics at concentrations previously validated for selecting transformants of other obligates or vehicle control beginning at four hpi. At 120 h post treatment, the cells were fixed, immunolabeled with antibody against the *O. tsutsugamushi* immunodominant outer membrane protein, TSA56 (56-kDa type-specific antigen), and qualitatively assessed by immunofluorescence microscopy for *Orientia* microcolony expansion in the cytosol. Chloramphenicol and rifampicin halted oriential growth while spectinomycin and gentamycin failed to do so at all concentrations tested (Fig. 3A). As a quantitative approach, qPCR was used to validate that 5 ug ml^-1^ chloramphenicol and 0.125 ug ml^-1^ rifampicin prevented *O. tsutsugamushi* replication (Fig. 3B). Next, we utilized the MTT assay to confirm that neither antibiotic impaired host cell metabolic activity even after 120 h of treatment (Fig. 3C, D). These data qualify chloramphenicol and rifampicin as potential selection agents for *O. tsutsugamushi*. It has been reported that acquisition of a single point mutation in *rpoB* renders obligate intracellular bacteria rifampicin-resistant (64, 65). Accordingly, even though we confirmed that our *O. tsutsugamushi* Ikeda str. does not carry this mutation via sequencing *rpoB*, we prioritized chloramphenicol as the selection agent.

**FIG 3.**
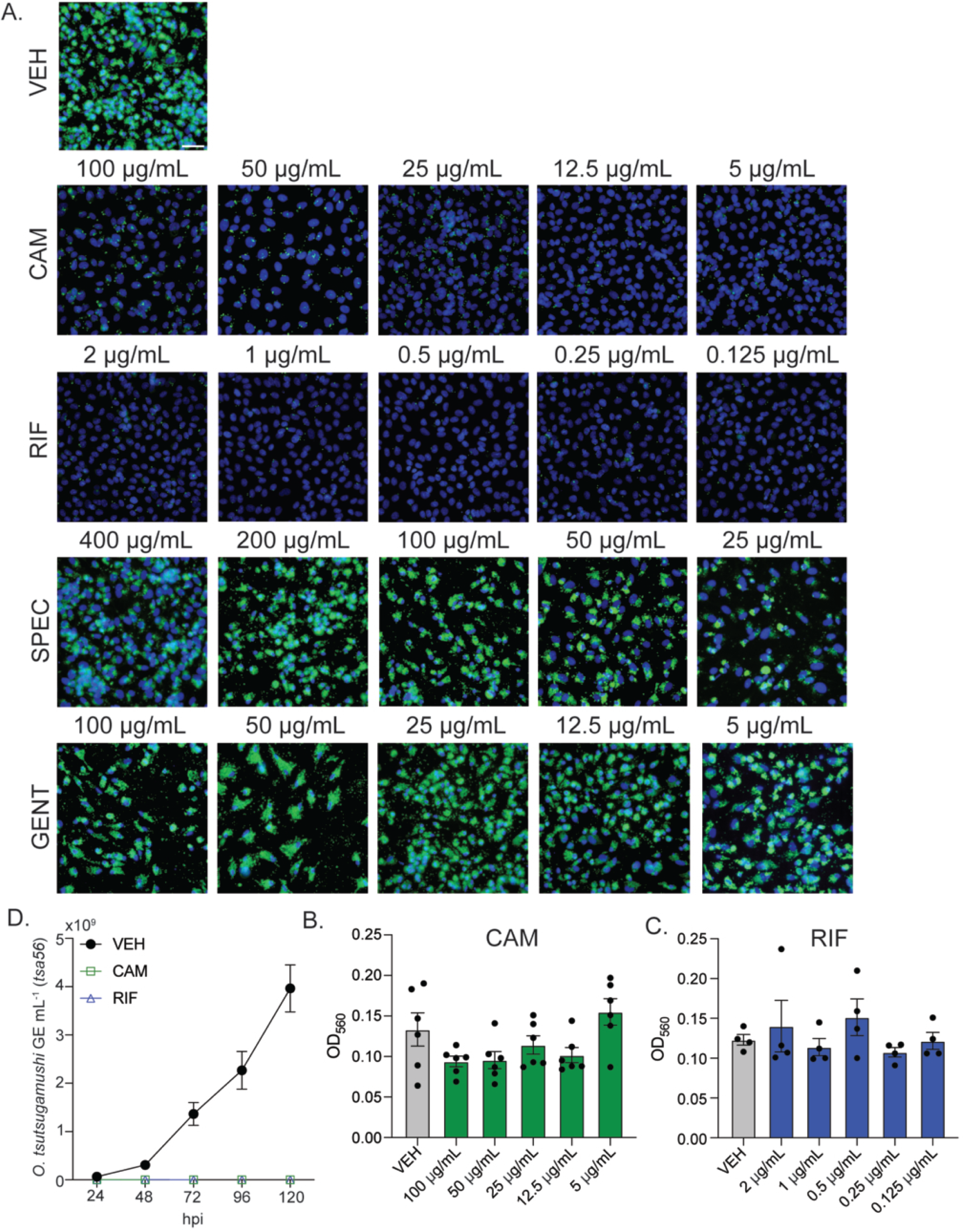
Assessment of antibiotic selection markers for *O. tsutsugamushi*. (A) EA.hy926 cells were infected with *O. tsutsugamushi* and treated at 4 hpi with vehicle control (VEH) or chloramphenicol (CAM), rifampicin (RIF), spectinomycin (SPEC), or gentamycin (GENT) at the indicated concentratinos. At 120 hpi, samples were fixed, immunolabeled with antibody against *O. tsutsugamushi* TSA56 (green), and stained with DAPI (blue) followed by immunofluorescence microscopy imaging with a 40X objective lens. Scale bar, 50 µm. (B) EA.hy926 cells were infected with *O. tsutsugamushi* and treated with CAM (5 µg mL^-1^), RIF (0.125 µg mL^-1^), or VEH at 4 hpi. qPCR was performed on total DNA isolated at 24, 48, 72, 96, and 120 hpi to determine GE/mL. Data are means ± SD and representative of three experiments with similar results. (C, D) An MTT assay was performed on EA.hy926 cells that had been treated with varying concentrations of CAM (C), RIF (D), or VEH (C, D) at 120 h. Data are means ± SEM and representative of 3–6 experiments with similar results. One-way ANOVA with Dunnett’s post hoc test was used to test for a significant difference compared to VEH. Statistically significant values are indicated.

### Generation of a plasmid for homologous recombination in *O. tsutsugamushi*

Due to the reductive evolution of the *O. tsutsugamushi* chromosome, it is presumed that most of its genes are essential. We devised a targeted homologous recombination strategy that would not delete or interrupt any annotated gene. Single copy *ank13* (OTT_RS04140) in Ikeda encodes Ank13, an ankyrin repeat-containing effector and nucleomodulin (25, 28). We constructed a plasmid (pJHOT::CM*ank13*) containing an *ank13* gene that is codon-modified (CM*ank13*) such that it is distinguishable from endogenous *ank13* but still conforms to the AT-richness of the *O. tsutsugamushi* chromosome (21). Arranged downstream from CM*ank13* are genes for the red fluorescent protein, mScarlet-I (66) and chloramphenicol acetyltransferase (CAT) (67) that have been codon-optimized for expression in *O. tsutsugamushi* and are under control of the *tsa22*-up and *tsa56*-down promoters, respectively (Fig. 4A). We previously established that these promoters are active throughout *O. tsutsugamushi* infection and drive high level expression (68). The mScarlet-I and CAT genes were placed tail-to-tail and separated by the synthetic DT5 terminator. The cassette was flanked upstream of CM*ank13* by sequence corresponding to Ikeda chromosome nucleotides 901,239–902,239, which included the predicted endogenous *ank13* promoter. The opposite end of the cassette, upstream from *tsa56*-down, was sequence corresponding to Ikeda chromosome nucleotides 897,766–899,766. For our homologous recombination strategy, we PCR amplified the cassette plus 1000 base pairs flanking either side and transform the purified 5292 bp product into *O. tsutsugamushi*. If successful, this would result in replacement of the 1473 bp *ank13* gene with the 3278 bp CM*ank13* cassette consisting of CM*ank13*, mScarlet-I, and CAT coding sequences (Fig. 4B).

**FIG 4.**
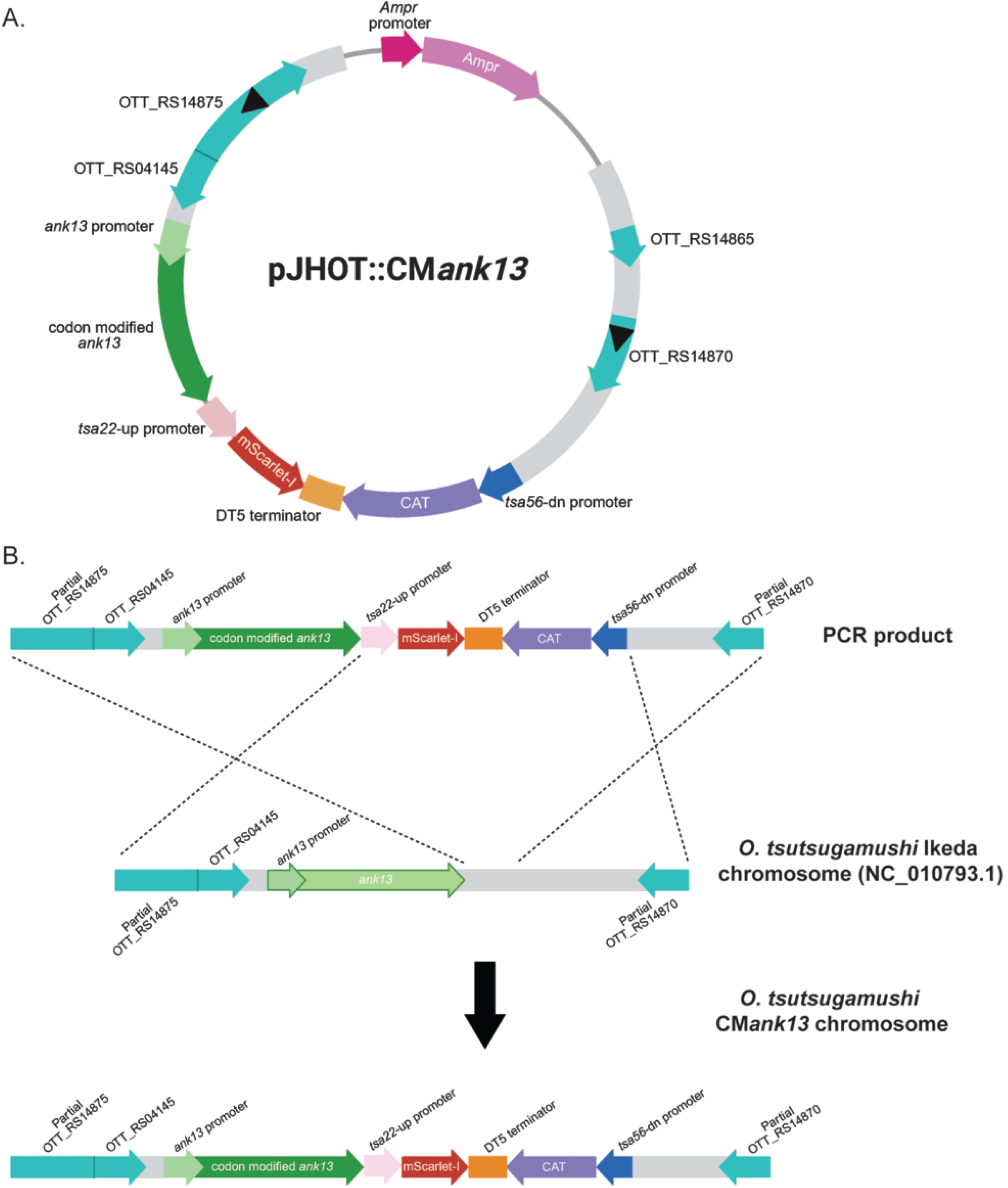
Plasmid and homologous recombination scheme for *O. tsutsugamushi*. (A) pJHOT::CM*ank13* plasmid map containing CM*ank13* (green) and the *ank13* native promoter (light green), mScarlet-I gene (red) under control of the *tsa22*-up promoter (pink), DT5 synthetic terminator (DT5; orange), chloramphenicol acetyltransferase (CAT) gene (purple) under control of the *tsa56*-down (*tsa56*-dn) promoter (blue). Regions corresponding to the *O. tsutsugamushi* str. Ikeda chromosome (NC_010793.1) that flank *ank13* are grey with genes denoted in turquoise. The ampicillin resistance gene (Ampr) and its promoter are colored pink and magenta, respectively. Black arrow heads denote the sequences targeted by primers for generating PCR product for transformation. (B) Homologous recombination scheme for insertion of the CM*ank13* cassette into the Ikeda chromosome. Dotted lines denote the predicted region for which homologous recombination between the Ikeda chromosome and PCR product will occur to generate the *O. tsutsugamushi* CM*ank13* chromosome.

### Comparison of CaCl_2_ and electroporation for transforming *O. tsutsugamushi* and on host cell integrity

The ability of *Orientia* to withstand common transformation methods had not been systematically evaluated. We compared CaCl_2_-based transformation, often employed for *Chlamydia* spp. (69, 70), and electroporation, which has been used to transform a variety of obligate intracellular bacteria (38, 39). The effects of each treatment on bacterial viability, host cell health, and DNA delivery were assessed. Figure 5A presents a schematic of the PCR-amplified fragment encompassing the CM*ank13* cassette and flanking regions that was used for transformation. Following CaCl_2_ treatment, *Orientia* viability decreased by approximately 50% and 75% in the absence and presence of PCR product, respectively, relative to untreated control bacteria (Fig. 5B, C). Electroporation killed ≥ 90% of the bacteria whether DNA was present or not. Following each transformation procedure, bacteria were added to EA.hy926 monolayers. Seventy-two h later, monolayer integrity was assessed by light microscopy. Electroporated *Orientia* caused substantial monolayer disruption compared to CaCl_2_-treated and control bacteria (Fig. 5D). Also, at 72 hpi, host cell-free *O. tsutsugamushi* organisms were recovered from the monolayers subjected to PCR and qPCR to detect the CM*ank13* cassette. pJHOt::CM*ank13* served as a positive control. PCR using the Mut0-F/R primer set (Fig. 5A) confirmed the presence of the cassette in bacteria that had undergone CaCl_2_ transformation or electroporation but not in untransformed or wild-type *O. tsutsugamushi* (Fig. 5E). qPCR using primer sets that targeted within the CAT or DT5 terminator sequences demonstrated that electroporation delivered DNA more efficiently than CaCl_2_-based transformation (Fig. 5F, G). Thus, although electroporation improves DNA uptake efficiency, its detrimental effect on both *O. tsutsugamushi* and host cell viability reduces the likelihood of recovering viable mutants. Conversely, whereas CaCl_2_ delivers DNA into *Orientia* less efficiently than electroporation, it is gentler on the bacteria and better preserves host cell integrity, making it the preferred method for transformation.

**FIG 5.**
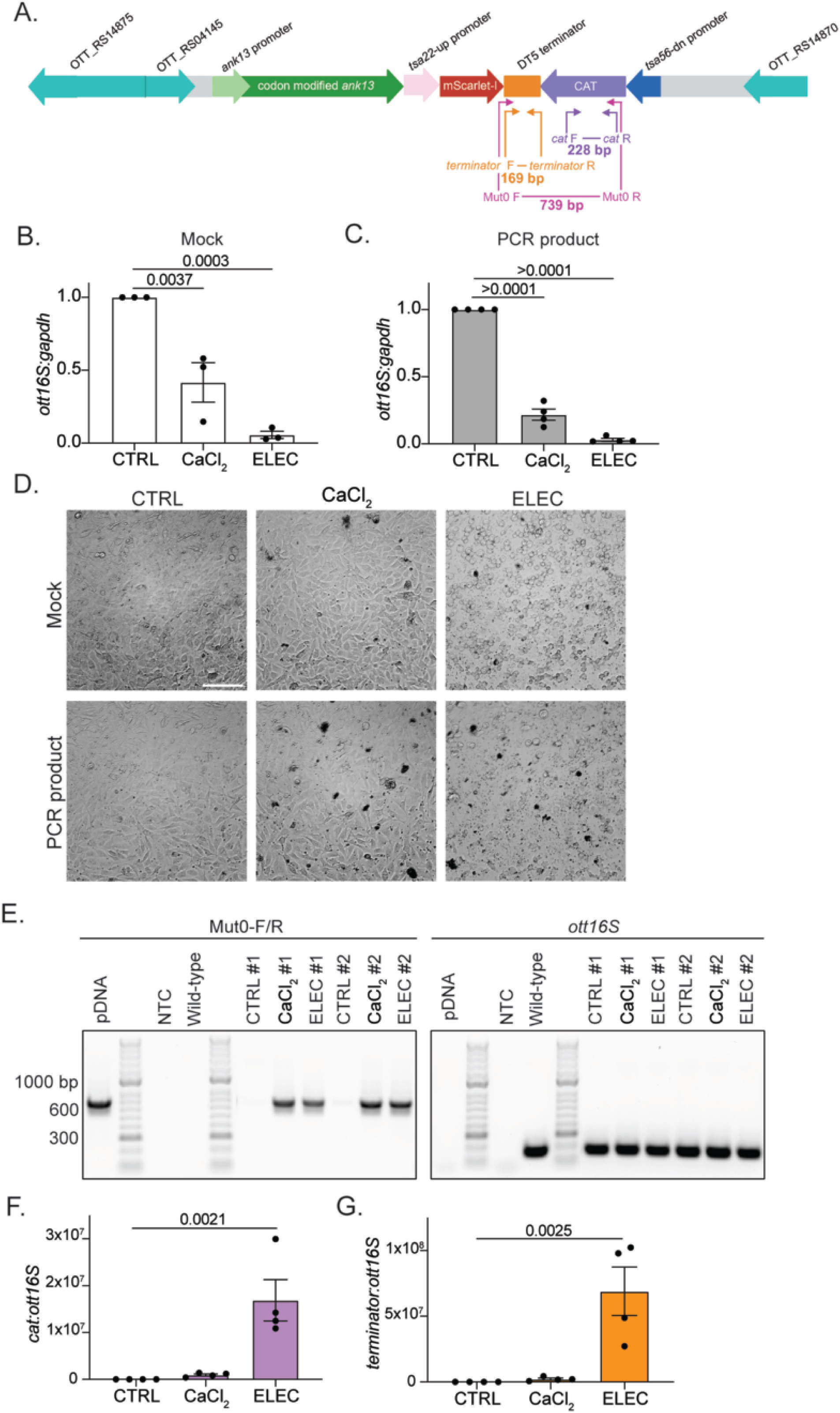
Comparing transformation methods for *O. tsutsugamushi*. (A) Schematic of the PCR-amplified fragment containing the CM*ank13* cassette and flanking regions that was used to transform *O. tsutsugamushi*. The annealing locations of and expected amplicon sizes for three different primer sets are indicated. (B, C) Assessment of transformation methods on *O. tsutsugamushi* viability. *O. tsutsugamushi* bacteria were incubated with CaCl_2_, subjected to electroporation (ELEC), or incubated at room temperature for 30 min (CTRL) without (Mock; B) or with 10 µg of PCR product (C) prior to being inoculated into naïve EA.hy926 cells. Total DNA isolated at 72 hpi was subjected to qPCR to determine the relative ratio of *O. tsutsugamushi 16s* (*ott16s*) to host *gapdh* as a measure of the bacterial load and indirect measure of bacterial viability. Data are means ± SEM. (D) Electroporated *O. tsutsugamushi* substantially disrupt host cell monolayer integrity. Phase microscopy was used to assess EA.hy926 monolayer integrity at 72 h post addition of *O. tsutsugamushi* organisms that had been subjected to CaCl_2_-mediated transformation or ELEC in the presence of PCR product. Images were acquired with a 40X objective lens. Scale bar, 50 µm. (E–G) Detection of the CM*ank13* cassette in *O. tsutsugamushi* post transformation. EA.hy926 cells were inoculated with *O. tsutsugamushi* organisms that had been transformed via CaCl_2_ or ELEC, or CTRL bacteria. Seventy-two h later, host cell-free bacteria were recovered. DNA was isolated and subjected to PCR using the Mut0-F/R primer set (A) and primers targeting *ott16S*. pDNA, pJHOt::CM*ank13*. Amplicons were resolved by agarose gel electrophoresis. NTC, no template control. Wild-type, DNA isolated from untransformed *O. tsutsugamushi*. The #1 and #2 designations refer to representative individual transformation experiments. qPCR was also performed using the *CAT*-F/R (A), Terminator-F/R (A), and *ott16S* primer sets and the *CAT*:*ott16S* and Terminator:*ott16S* ratios quantitated. Data are representative of 3-4 experiments with similar results. One-way ANOVA with Dunnett’s post hoc test was used to test for a significant difference compared to CTRL. Statistically significant values are indicated.

### Generation of transgenic *O. tsutsugamushi* by allelic exchange

Having established optimal growth conditions, antibiotic selection parameters, and transformation methodology, we developed a preliminary workflow for transformation, outgrowth, and confirmation of allelic exchange leading to generation of the *O. tsutsugamushi* CM*ank13*. CaCl_2_-based transformation was performed with 10^8^–10^9^ host cell-free *O. tsutsugamushi* bacteria, as confirmed by GE quantification, and 10 µg of PCR product. Transformed bacteria were introduced into a 75-cm^2^ cell flask containing 100% confluent naïve EA.hy926 cells and allowed to replicate for 72 hpi at which time cytopathic effects indicating an active infection were observed. Chloramphenicol was added to a final concentration of 5 µg mL^-1^. Media with chloramphenicol was replaced every 72–120 h. Beginning at 72 h post chloramphenicol addition and every 14 days thereafter, to screen for the presence of *O. tsutsugamushi* CM*ank13* mutants, infected cells were trypsinized and processed for bacterial isolation and PCR analysis. Aliquots were also seeded onto coverslips and fixed for immunofluorescence microscopy to detect colocalization of endogenous mScarlet-I fluorescence with *O. tsutsugamushi* TSA56 immunosignal.

Figure 6A presents schematics of the relevant region of the *O. tsutsugamushi* chromosome with and without the integrated CM*ank13* cassette. Binding sites of PCR primer sets along with the predicted amplicon sizes are indicated. PCR using the Mut0-F/R, Mut1-F/R, and Mut2-F/R primer pairs, which bind within the CM*ank13* cassette (Fig. 6A), yielded products of the expected size for *O. tsutsugamushi* CM*ank13* but not wild-type *O. tsutsugamushi* (Fig. 6B). PCR with the Mut3-F/R primer set, which targets the flanking chromosomal regions outside the cassette (Fig. 6A), yielded a 5292 bp amplicon for transgenic *O. tsutsugamushi* compared to a 3487 bp amplicon in wild-type *Orientia* (Fig. 6C). The Mut4-F and Mut4-R primers bind within the cassette CAT gene and flanking chromosomal sequence, respectively (Fig. 6A), and therefore would only yield product if the CM*ank13* cassette had properly integrated at the predicted chromosomal location. PCR with Mut4-F/R generated a product of the expected 2.8 kB size for *O. tsutsugamushi* CM*ank13* but not wild-type *Orientia* (Fig. 6D). Gel purification and nanopore sequencing of the Mut3-F/R and Mut4-F/R amplicons verified the sequence integrity of each. Sequencing of the three smaller Mut3-F/R products generated from transgenic *O. tsutsugamushi* confirmed that they were the result of non-specific amplification. At 14 days post transformation, *O. tsutsugamushi* CM*ank13* was isolated, total DNA collected, and PCR with *ank13*-F/R and CM*ank13*-F/R primer pairs (Fig. 6A) was performed. Although wild-type *ank13* was present in both wild-type and mutant *O. tsutsugamushi* samples, CM*ank13*-F/R primers produced an amplicon only in *O. tsutsugamushi* CM*ank13* (Fig. 6E). Anti-TSA56 immunolabeled mScarlet-I positive perinuclear *O. tsutsugamushi* microcolonies (Fig. 6F). This workflow was successfully performed nine times, each yielding *O. tsutsugamushi* CM*ank13* mutants as confirmed by PCR and sometimes also immunofluorescence microscopy. Unfortunately, we were unable to maintain *O. tsutsugamushi* CM*ank13* populations beyond approximately 28 days (Fig. 6G). Results in Figure 6G are representative for all nine attempts. Nonetheless, we had successfully transformed and achieved allelic exchange in *O. tsutsugamushi*.

**FIG 6.**
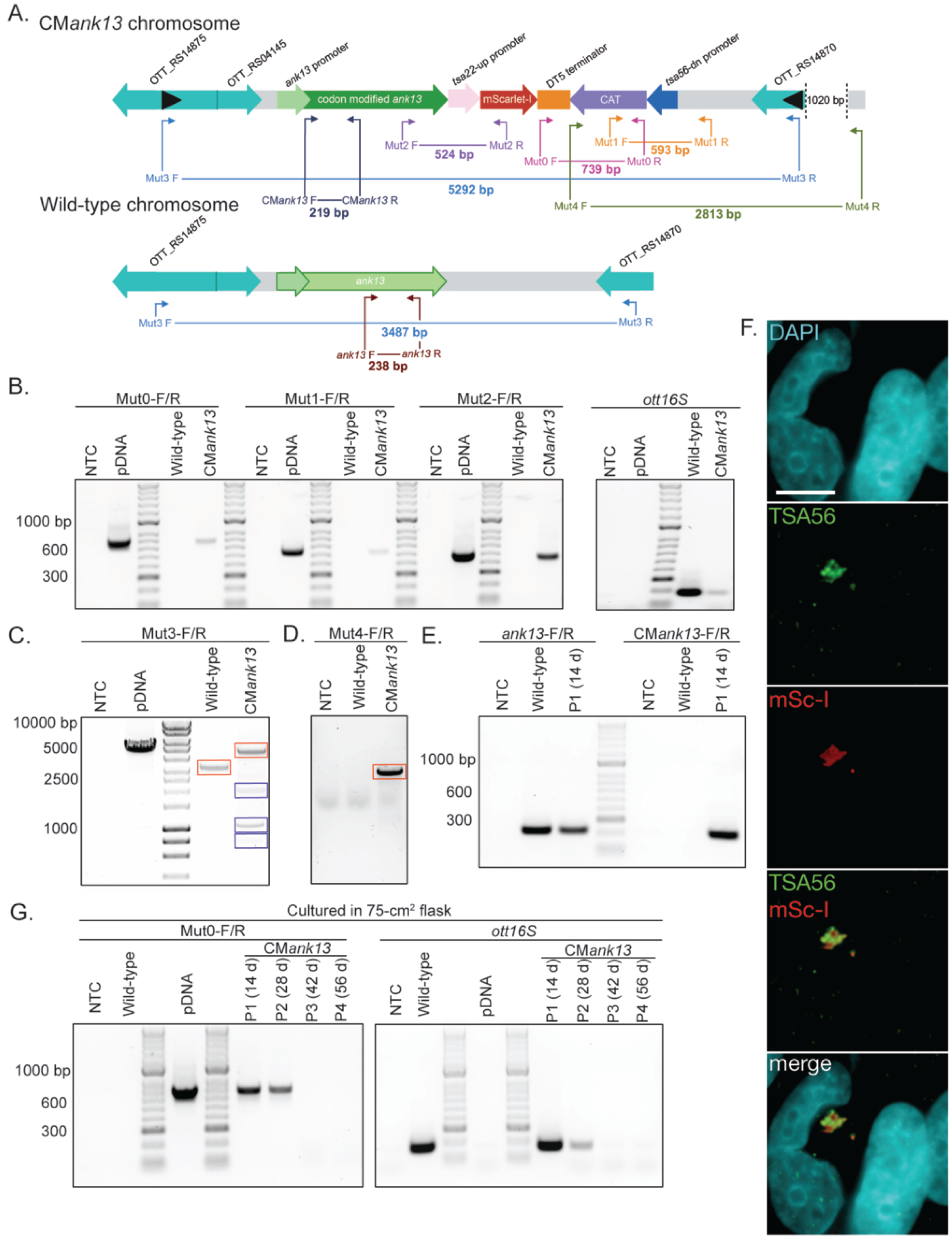
Confirmation of CM*ank13* allelic exchange in *O. tsutsugamushi*. (A) Schematics of the *O. tsutsugamushi* str. Ikeda chromosome with and without the integrated CM*ank13* cassette integrated. Annealing locations of and expected amplicon sizes for the indicated primer sets are provided. (B–D) PCR confirmation of allelic exchange. Transgenic *O. tsutsugamushi* bacteria that had been transformed with PCR product containing the CM*ank13* cassette and cultivated in EA.hy926 cells for two weeks were recovered. (B) DNA was isolated and subjected to PCR using the indicated primer sets. Amplicons were resolved by agarose gel electrophoresis. pJHOT::CM*ank13* plasmid DNA (pDNA) served as a positive control while untransformed *O. tsutsugamushi* (Wild-type) and no template (NTC) were negative controls for amplification. The *ott16S* primer served as a positive control to amplify *O. tsutsugamushi* DNA. bp, base pair. Amplicons generated using the Mut3-F/R (C) and Mut4-F/R (D) primer sets were subjected to nanopore sequencing. Amplicons outlined in red represent sequences that produced the expected product while those outlined in blue denote products resulting from non-specific amplification. (E) Total DNA isolated from EA.hy926 cells containing *O. tsutsugamushi* CM*ank13* was subjected to PCR using the indicated primer sets. Amplicons were resolved by agarose gel electrophoresis. Wild-type *O. tsutsugamushi* was used as a positive control for amplifying wild-type *ank13* and a negative control for CM*ank13* amplification. NTC was a negative control. (F) EA.hy926 cells harboring mutagenized *O. tsutsugamushi* were seeded on a coverslip followed by paraformaldehyde fixation at 72 h, immunolabeling for *O. tsutsugamushi* TSA56 (green), and staining with DAPI (blue). Immunofluorescent microscopy was used to visualize native mScarlet-I (mSc-I; red) fluorescence with TSA56 and DAPI signals. Images were acquired with a 100X oil immersion objective lens. Scale bar, 10 µm. (G) *O. tsutsugamushi* CM*ank13* bacteria were culture in 75-cm^2^ flasks containing EA.hy926 cells. DNA isolated from host cell-free *O. tsutsugamushi* CM*ank13* bacteria at 14 d, 28 d, 42 d, and 56 d was subjected to PCR with Mut0-F/R and *ott16S* primers.

### Workflow optimization using live-cell fluorescence microscopy facilitates recovery and maintenance of transgenic *O. tsutsugamushi* CM*ank13*

Given our inability to sustain *O. tsutsugamushi* CM*ank13* bacteria in EA.hy926 cell monolayers in 75-cm^2^ flasks beyond four weeks, we refined our workflow (Fig. 7). CaCl_2_-mediated transformation, inoculation into EA.hy926 cells in a 75-cm^2^ flask, and cultivation in the absence of selection for 48–72 h followed by the addition of chloramphenicol was performed as above. Twenty-four h post chloramphenicol addition, the culture was treated with trypsin, and the cells were transferred evenly among the wells of a 96-well glass-bottom plate. Cultivation was continued under chloramphenicol selection, with media changes every 72 h, for two weeks after which the wells were imaged by live-cell microscopy. Ten fields of view per well were imaged by phase contrast and fluorescence microscopy for a total of 960 merged images per plate. The images were screened to denote wells with cells harboring fluorescent *O. tsutsugamushi* organisms. Over multiple experiments, 10–13 wells of a 96-well plate tended to contain transgenic bacteria. After three consecutive weeks in the presence of chloramphenicol the host cells displayed irregular growth dynamics and increased vacuolation, suggesting long-term chloramphenicol treatment has negative impacts on host cell health. Accordingly, we then alternated media with or without chloramphenicol every 72–120 h. After two weeks, each culture of interest was transferred to a well of a 24-well glass-bottom plate and cultivation under antibiotic selection was continued for an additional five days. Imaging this plate (10 fields of view per well) revealed that the cultures of interest retained mScarlet-I positive bacteria. The plate was subsequently imaged every one to two weeks to confirm the presence of transgenic *O. tsutsugamushi*. Fresh EA.hy926 cells were added if the infection progressed to lyse ≥ 50% of the cells. At week 10, each well was split to two duplicate wells that were maintained in parallel with or without chloramphenicol. At week 12, Hoechst dye was added to wells for each condition. Live-cell imaging revealed that the transgenic bacteria exhibited mScarlet-I signal colocalizing with Hoechst-stained nucleoids, while wild-type control *O. tsutsugamushi* were only Hoechst-positive (Fig. 8A). Importantly, background autofluorescence signal did not colocalize with *Orientia* nucleoids for any condition. At week 13 (88 d), DNA was isolated from cultures containing chloramphenicol. PCR with primer sets specific for wild-type *ank13* or CM*ank13* confirmed that the mScarlet-I positive bacteria were in fact *O. tsutsugamushi* CM*ank13* (Fig. 8B). An extremely faint wild-type *ank13* amplicon was detectable in wild-type *Orientia* and *O. tsutsugamushi* CM*ank1*3, indicating that the wild-type bacteria had been nearly cured from the transgenic *Orientia* culture.

**FIG 7.**
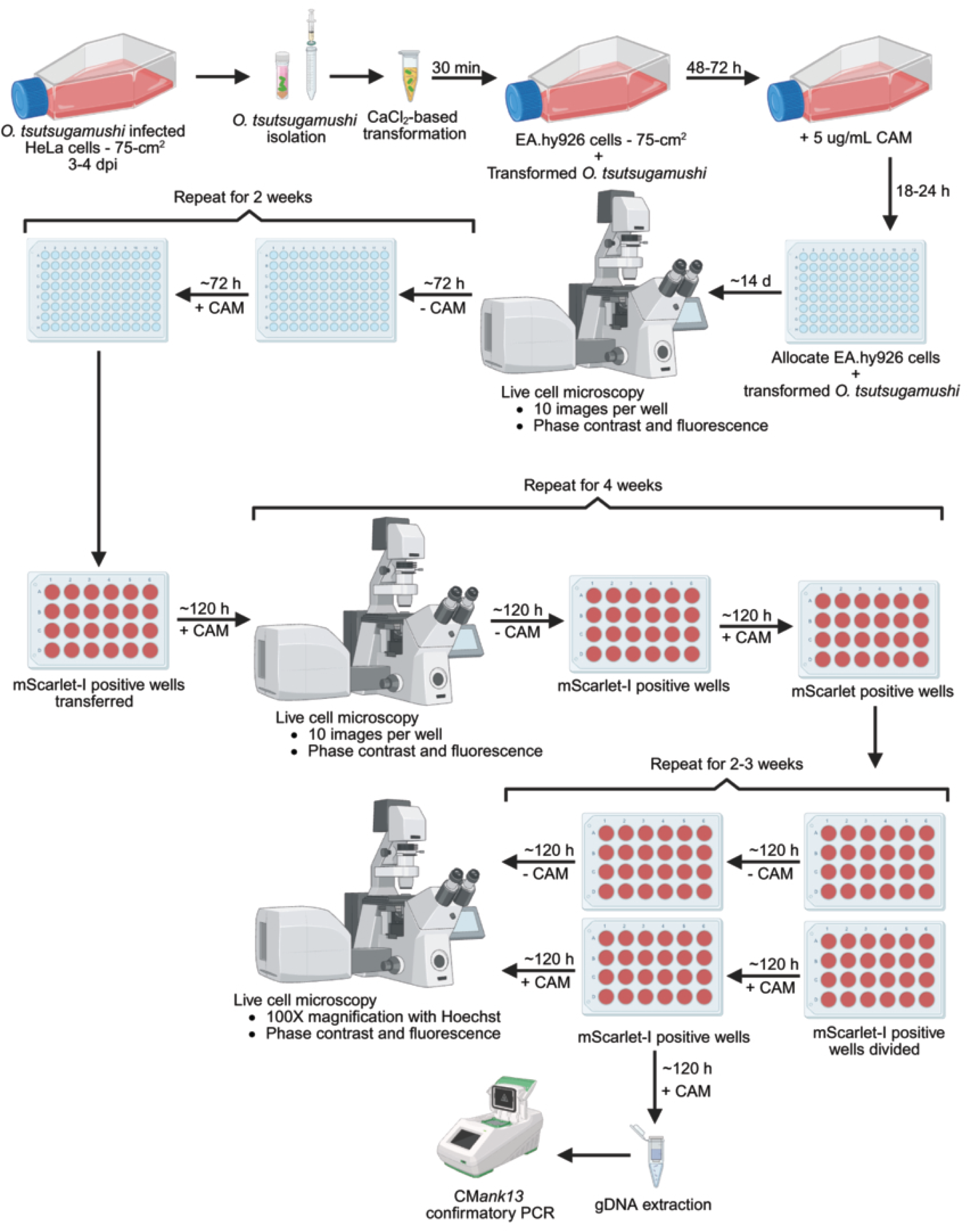
Workflow for transforming and recovering transgenic *O. tsutsugamushi* CM*ank13* organisms. *O. tsutsugamushi* (10^8^–10^9^) bacteria were isolated and subjected to CaCl_2_-based transformation with 10 µg of CM*ank13* PCR product. Transformed bacteria were inoculated onto an EA.hy926 cell monolayer in a 75-cm^2^ culture flask and incubated for 48-72 h before addition of 5 µg mL^-1^ chloramphenicol (CAM). At 18-24 h post chloramphenicol addition, cultures were treated with trypsin, allocated evenly among the wells of a 96-well glass-bottom plate, and incubated for 14 d. Next, ten phase and fluorescent micrographs per well were acquired by live cell microscopy. mScarlet-I positive cultures were maintained for two weeks during which media was alternated with or without CAM every 72 h. After two weeks, mScarlet-I positive cultures in individual wells were treated with trypsin and transferred to individual wells of a 24-well glass-bottom plate. Cultures were maintained for four weeks during which media with or without CAM was alternated every 120 h. During this four-week period, the wells were monitored by live cell microscopy every 1-2 weeks. At 10 weeks post transformation, each mScarlet-I positive culture was split and maintained with or without CAM in parallel. Representative cultures of EA.hy926 cells harboring *O. tsutsugamushi* CM*ank13* was were subjected to confirmatory PCR or imaged by live cell microscopy with Hoechst staining.

**FIG 8.**
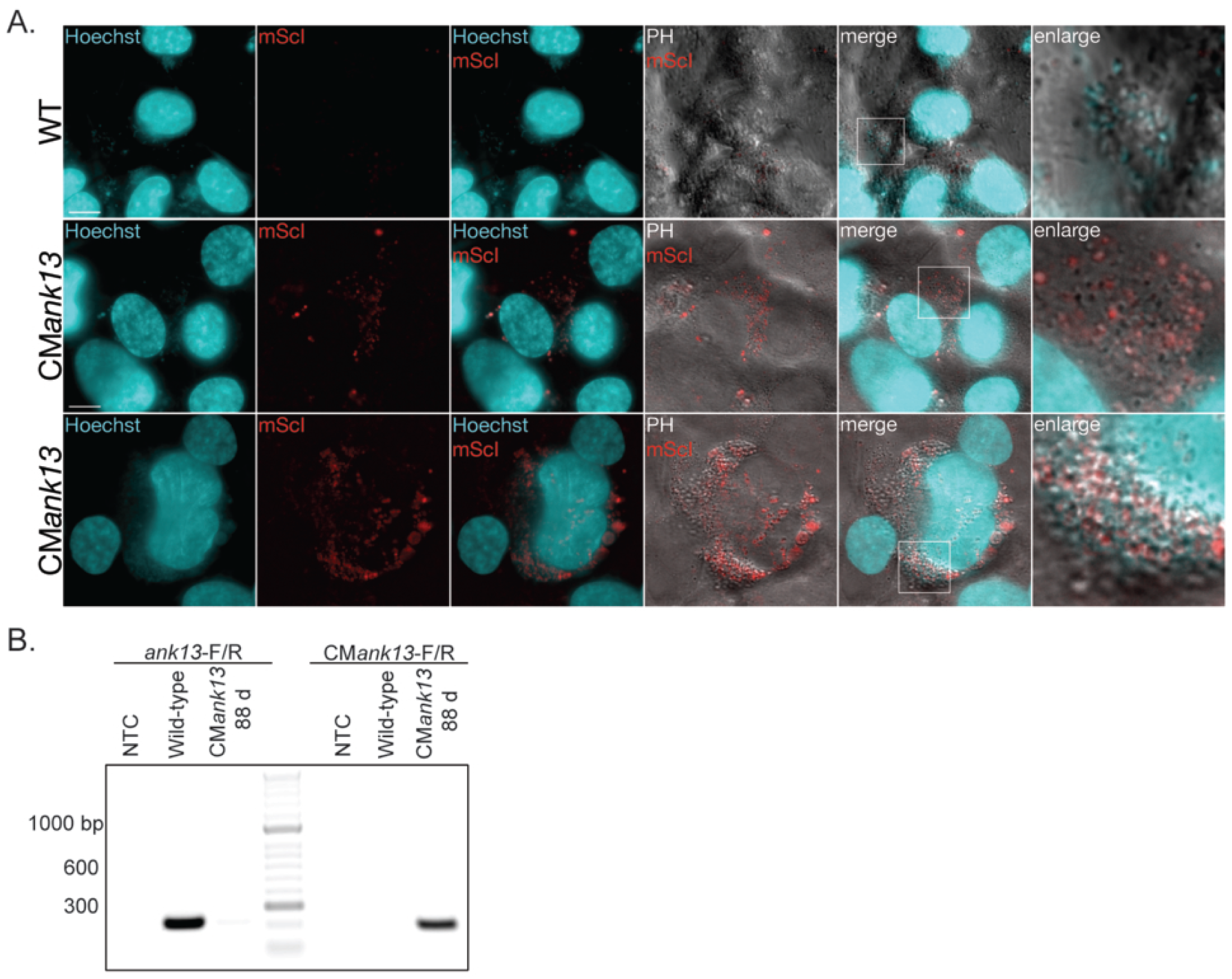
Confirmation and maintenance of *O. tsutsugamushi* CM*ank13* organisms in culture. (A) EA.hy926 cells harboring wild-type of CM*ank13 O. tsutsugamushi* in 24-well glass-bottom plates were stained with Hoechst followed by live cell imaging. Images were acquired with a 100X oil immersion objective lens. Scale bar, 10 µm. (B) Total DNA isolated from EA.hy926 cells infected with *O. tsutsugamushi* CM*ank13* bacteria were collected and subjected to PCR using the indicated primer sets. Amplicons were resolved by agarose gel electrophoresis. Untransformed *O. tsutsugamushi* (Wild-type) was used as a positive control for wild-type *ank13* amplifcation and a negative control for *CMank13* amplification. No template (NTC) was a negative control for amplification in general. The *ott16S* primer served as a positive control to amplify *O. tsutsugamushi* DNA. bp, base pair.

## DISCUSSION

The lack of tools for genetically manipulating *O. tsutsugamushi* has limited understanding of the fundamental bacterial biology and host-microbe interactions that shape scrub typhus pathogenesis. We hope that reporting our investigations that enabled us to transform *O. tsutsugamushi*, achieve and verify allelic exchange, visualize transgenic organisms, and maintain them in culture will accelerate development of additional genetic tools for this understudied pathogen. We selected the Ikeda for this study due to the severity of infection that it causes in humans and laboratory mice (17, 55–58), its genome being well annotated (16, 21), and our extensive experience in working with it (24–32, 36, 37, 71–73). That said, given that virulence severity varies among strains (74), it will be crucial to confirm that the approaches described herein will be applicable to other clinical strains, particularly Karp- and Gilliam-related strains, which account for approximately 50% and 25% of scrub typhus infections, respectively (17, 19, 74).

Although CaCl_2_ delivers DNA into *O. tsutsugamushi* less efficiently than electroporation, incubating electroporated *Orientia* organisms with host cells results in considerably more cytoxicity versus CaCl_2_-transformed bacteria. This is likely because electroporation-mediated damage to the *Orientia* cell wall releases damage-associated molecular patterns that induce host cell death pathways. We infer that CaCl_2_ results in less bacterial cell death, which is important considering that a ratio of greater than one viable *O. tsutsugamushi* organism per host cell is optimal for a productive infection. Thus, between the two transformation methods evaluated, CaCl_2_-mediated transformation is preferable. Of the three cell lines assessed, EA.hy926 proved optimal for enabling high *Orientia* loads and preventing host cell overgrowth presumably due to contact-dependent inhibition (75). These properties translated to EA.hy926 cells being excellent for supporting the outgrowth of rare transgenic *O. tsutsugamushi* CM*ank13* organisms. Importantly, cells harboring mScarlet-I positive transgenic bacteria could easily be identified when grown in 96- or 24-well glass bottom plates and visualized by live-cell fluorescence microscopy, allowing us to select and expand these cultures.

*O. tsutsugamushi* str. Ikeda did not form plaques in EA.hy926 cells after 14 days. Our protocol was based on Hanson’s (61), which showed that *O. tsutsugamushi* strains Karp, Kato, and Gilliam formed plaques in the murine fibroblast line, C3H/10T1/2 clone 8 within a similar observation period to ours. These cells, like EA.hy926 cells, exhibit contact-dependent inhibition. Gamma irradiation or treatment with daunomycin or cycloheximide is not needed for C3H/10T1/2 clone 8 cells to support *O. tsutsugamushi* plaque formation (61). Whether EA.hy926 cells grow faster than C3H/10T1/2 clone 8 cells, which would conceivably allow them to fill zones of clearing and hence prevent plaque formation, is an untested possibility. In Hanson’s protocol, *Orientia* inocula were purified from chicken egg cultures and the initial MOI was not determined (61). The MOI range that we employed with bacteria isolated from EA.hy926 cells could have been comparatively much lower. If so, then a higher *Orientia* inoculum load or longer cultivation period may be needed for plaque formation. Another notable difference between the protocols is the serum supplementation employed. We provided *Orientia* infected EA.hy926 cells with 10% or 1% FBS. Hanson found that 5% FBS + 5% chicken serum increased *Orientia* plaque formation three-fold and increased plaque size 1.6-fold versus 10% FBS (61). Now that we have a reliable transformation protocol for *O. tsutsugamushi*, it will be worth testing if EA.hy926 or C3H/10T1/2 clone 8 cells supplemented with 5% FBS + 5% chicken serum and potentially inoculated with higher MOIs than we employed in this study will enable plaque formation and hence recovery of isogenic mutants.

Chromosomal insertion of the CM*ank13* cassette was confirmed by PCR and nanopore sequencing. Allelic exchange was achieved by including 1000 bp flanking either side of the cassette, which can serve as a guideline for future allelic exchange approaches for *O. tsutsugamushi*. It will be worth evaluating if smaller flanking regions can still mediate homologous recombination as a means for decreasing overall insert size to increase transformation efficiency. Codon-optimized CAT expression driven by the *tsa56*-down promoter could be inferred by the ability of transformants to grow in the presence of chloramphenicol while expression of codon-optimized mScarlet-I from the *tsa22*-up promoter was confirmed by visualization of fluorescent bacteria. The ability of transgenic *O. tsutsugamushi* to be selected and maintain mScarlet-I expression over at least 13 consecutive weeks validates the efficacy of these promoters for driving transgene expression in the bacterium. Based on a promoter activity assay in *Escherichia coli*, *tsa22*-up exhibited the most robust activity followed by *tsa56*-down (68). Two other promoters evaluated in that study, *tsa22*-down and *tsa56*-up, displayed comparable activity to that of *tsa56*-down and could therefore be considered for driving transgene expression in *Orientia*. Moreover, there are likely additional promoters that could drive constitutive gene expression more robustly than those evaluated thus far and therefore improve resistance and/or reporter gene expression. Indeed, very little is known regarding how *O. tsutsugamushi* gene expression varies throughout the infection cycle, during residence in different host cell types, or between its recently identified distinct morphological forms (68, 76). RNAseq and RT-qPCR analyses examining these conditions would greatly augment selection of additional *O. tsutsugamushi* gene promoters for evaluation.

Of the antibiotics evaluated that inhibited *O. tsutsugamushi* growth, chloramphenicol was prioritized over rifampicin due to the potential of RpoB to become resistant to rifampicin following a single nucleotide mutation (64, 65). While chloramphenicol has been used to treat scrub typhus and infections caused by other obligate intracellular bacteria (46, 65, 77–82), it is typically advised against due to its potential life-threatening side effects such as aplastic anemia (83). Importantly, chloramphenicol-resistant *O. tsutsugamushi* would be susceptible to the first-line drug of choice, doxycycline, and other clinically employed antibiotics including azithromycin, an excellent alternative to doxycycline (63). Chloramphenicol can inhibit mitochondrial protein synthesis resulting in stress to the organelle and decreased ATP synthesis (84). Although the MTT assay indicated that EA.hy926 cells did not exhibit mitochondrial stress after cultivation in five μg ml^-1^ chloramphenicol for five days, we cannot rule out that extended cultivation in its presence would induce at least some mitochondrial stress. If so, this could compromise the fitness of even chloramphenicol-resistant transgenic *Orientia* due to the bacterium’s interplay with mitochondria during infection (72, 73). Of note, wild-type *O. tsutsugamushi* is abolished by day 42 and *O. tsutsugamushi* CM*ank13* organisms can be maintained in the absence of chloramphenicol. Theoretically, this or any other chromosomal-insertion mutant ought to be maintainable in the absence of selection, which would eliminate the concern over chloramphenicol-induced mitochondrial stress. Nonetheless, it will be prudent to pursue alternative agents for selecting transgenic *O. tsutsugamushi* especially ones that are 100% clinically irrelevant for scrub typhus. Also, having more than one selection agent would pave the way for knock out-complementation approaches and fulfillment of Molecular Koch’s postulates.

Successful plasmid systems in other obligates require shuttle vectors derived from endogenous plasmids (38), which *Orientia* lacks (16, 21–23, 41). Plasmids have been found in *Rickettsia* spp. and appear to have originated from *Rickettsia*/*Orientia* chromosomes (38, 85–89). As noted by Fisher and Beare, this suggests that rickettsial shuttle vectors might be compatible with *Orientia* (38). Moreover, an origin of replication sequence has been identified in *O. tsutsugamushi* (20, 21), suggesting that it might be able to support an autonomously replicating plasmid. Applying this information to determine if *Orientia* can be transformed with and support a plasmid will be a key future endeavor.

In summary, we established a pipeline for transforming and achieving allelic exchange in *O. tsutsugamushi*. This work is a starting point for refining the methods reported herein and establishing more genetic tools for determining *O. tsutsugamushi* gene function, improving host-pathogen interaction studies, and developing attenuated mutants.

## MATERIALS AND METHODS

### Cultivation of cell lines and *O. tsutsugamushi* infections

Uninfected EA.hy926 human umbilical vein cells (CRL-2922; American Type Culture Collection [ATCC]), HeLa human cervical epithelial cells (CCL-2; ATCC), Vero green African monkey kidney cells (CCL-81; ATCC) were maintained as done previously (25, 60). HeLa cells infected with *O. tsutsugamushi* str. Ikeda were maintained as described (25). To isolate *O. tsutsugamushi*, infected host cells were collected by trypsinization and pelleted at 1000 x *g* for 3 min after which the supernatant was removed. The pellet was resuspended in 1X phosphate-buffered saline (PBS; 1.05mM KH_2_PO_4_, 155mM NaCl, 2.96mM Na_2_HPO_4_ [pH 7.4]) and combined with 0.7 mm zirconia beads (Biospec Products). The host cells were disrupted for 20 sec using a FastPrep-24 sample preparation system (MP Biomedicals) to release intracellular bacteria. Host cell debris was removed by filtration through a 2 µm Whatman Puradisc glass microfiber syringe filter (Cytiva). The resulting host cell-free *O. tsutsugamushi* was utilized for continuous cultivation in HeLa cells or infection studies (25). Infections were synchronized at 4 hpi by removing inocula and replacing with fresh complete Dulbecco’s modified Eagle’s medium (DMEM) with l-glutamine, 4.5 g d-glucose and 100 mg sodium pyruvate (Gibco) supplemented with 1% or 10% (vol/vol) heat-inactivated FBS (Gemini Bioproducts), 1X minimal essential medium containing non-essential amino acids (Gibco) and 15 mM HEPES (Gibco) for EA.hy926 and Vero cells or Roswell Park Memorial Institute medium (RPMI) with l-glutamine (Gibco) supplemented with 1% or 10% (vol/vol) heat-inactivated FBS for HeLa cells. MOI was assessed by immunofluorescence microscopy of coverslips of infected cells as described below. Experiments were verified for achieving a MOI of 5-10 in at least 95% of the host cells unless otherwise specified.

### GE quantification

Lysates for GE were collected by trypsinization followed by dilution in H_2_O and incubation at 100°C for 15 min as described previously (25). At the time of analysis, samples were thawed, vortexed, and GE quantified by qPCR. *O. tsutsugamushi* str. Ikeda *tsa56* gene (OTT_RS04590) (25) was amplified using gene-specific primers (Table 1) with PerfeCTa SYBR Green Fastmix (Quantbio), and a CFX384 Real Time PCR Detection System (Bio-Rad Laboratories). Thermal cycling conditions were 95°C for 30s followed by 40 cycles of 95°C for 10s to 54°C for 10s, and a 65°C-to-95°C melt curve. A standard curve was prepared using the plasmid, pCR4.0::*Ot_tsa56* (24). *O. tsutsugamushi* GE mL^-1^ was calculated based on the standard curve as done previously (25).

**Table 1.**
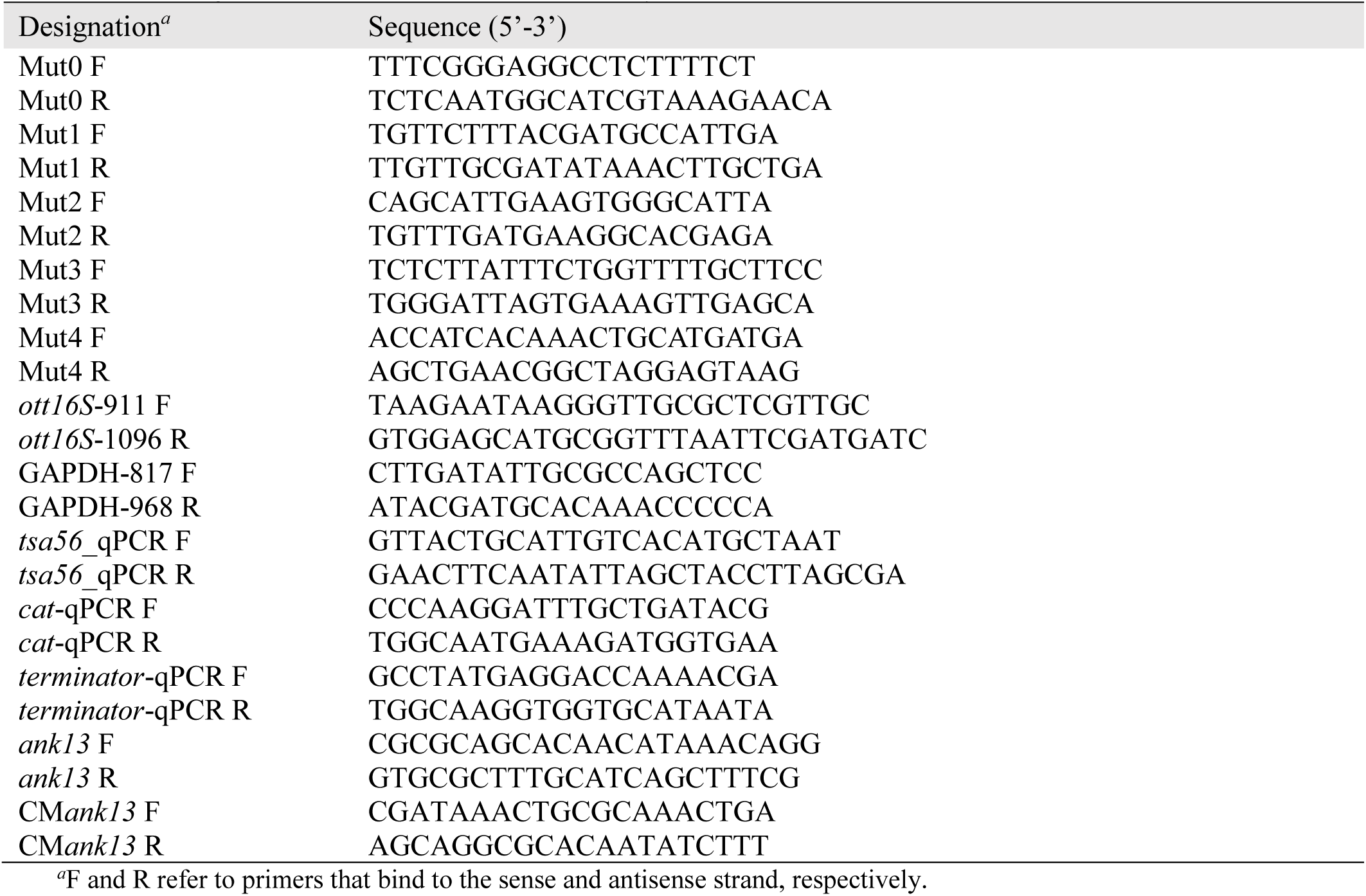
Oligonucleotides utilized in this study.

### Host cell proliferation assay

HeLa, EA.hy926, and Vero cells were seeded to achieve 60-70% confluency. Prior to synchronous infection with *O. tsutsugamushi*, cells were labeled using the CellTrace Far Red Proliferation kit (ThermoFisher Scientific) following the manufacturer’s protocol. Briefly, after removing the medium, 1 µM of CellTrace Far Red dye diluted in pre-warmed 1X PBS was added, and the cells were incubated at 35°C in a humidified incubator with 5% CO_2_ for 20 min. The dye solution was removed and cells washed twice with complete media. Cells were incubated in complete media at 35°C with 5% CO_2_ for 10 min prior to *O. tsutsugamushi* infection. At the time of analysis, cells were dislodged with 0.526 mM EDTA (Versene) solution and the fluorescence intensity of CellTrace Far Red dye in 10,000 individual cells was quantified using a Becton Dickinson (BD) FACSMelody equipped with BDFACSChorus 1.3.3 software (BD Biosciences). Post data-acquisition analyses were performed using FlowJo software version 10.8.1 (BD Biosciences) with population gates isolating single cell populations (25).

### Plaque assays

The protocol employed was comparable to that by Hanson with minor modifications (61). HeLa or EA.hy926 cells were seeded to achieve 100% confluency in six-well plates. Medium was removed and host cells were washed with 1X PBS prior to inoculation with 100 µL of *O. tsutsugamushi* isolated as described above to achieve an MOI of 30. Infected cells were incubated at 35°C with 5% CO_2_ and agitation for 15 min followed by the addition of an overlay composed of complete media with 1% or 10% FBS, with or without 100 mg mL^-1^ cycloheximide, and 0.5% SeaPlaque agarose (Lonza). Plates were incubated at 35°C with 5% CO_2_, visualized daily, and stained with neutral red or MTT at 14 dpi. Neutral red staining was achieved by adding 1.5 mL of 0.03% neutral red stain diluted in serum free DMEM to the initial overlay and incubating at 35°C with 5% CO_2_ overnight. MTT staining was performed by adding 1 mL of 0.5 mg mL^-1^ MTT stain diluted in 1X PBS to the initial overlay and incubating at 35°C with 5% CO_2_ for 4 h.

### Microscopy

EA.hy926 cells were seeded in 24-well plates with 12-mm round glass coverslips (Electron Microscopy Sciences) and synchronously infected with *O. tsutsugamushi*. At the time of collection or at 4 hpi for MOI quantification, cells were fixed in 100% methanol for 5 min followed by successive incubations with rabbit-anti TSA56 (30) (1:1,000) for 1 h in 1X PBS containing 5% (vol/vol) BSA and Alexa Fluor 488-conjugated goat anti-rabbit IgG (Invitrogen [A-11034]; 1:1000) plus 0.1 mg mL^-1^ 4′,6′-diamidino-2-phenylindole (DAPI; Invitrogen [D1306]). The coverslips were mounted using ProLong Gold antifade reagent (Invitrogen). Immunofluorescence and phase images were acquired with a TCS SP8 microscope (Leica Microsystems). For live cell microscopy, 96- or 24-well glass-bottom plates harboring EA.hy926 infected with *O. tsutsugamushi* CM*ank13* transformants were imaged in an Oko-Touch CO_2_-heated stage incubator at 35°C with 5% CO_2_. At 2 weeks post transformation and every 1-2 weeks thereafter, 10 phase and fluorescent micrographs per well were acquired at 20X magnification using a TCS SP8 microscope. At 12 weeks post transformation, cultures were stained with Hoechst stain (Invitrogen) and representative images of *O. tsutsugamushi* CM*ank13* transformants or wild-type *O. tsutsugamushi* were acquired at 100X. Micrographs were processed and analyzed using Fiji, (version 2.16.0/1.54g).

### Antibiotic treatments

EA.hy926 cells seeded in 24-well glass bottom plates (Sigma-Aldrich) were synchronously infected with *O. tsutsugamushi*. CAM (100–5 µg mL^-1^), RIF (2–0.125 µg mL^-1^), SPEC (400–25 µg mL^-1^), or GENT (100–5 µg mL^-1^) diluted in complete DMEM were added to infected cells at 4 hpi, and cultivation continued for 120 h. The bacterial load was qualitatively assessed using immunofluorescence microscopy as described above. For growth curve analyses, *O. tsutsugamushi* GE were quantified in CAM (5 µg mL^-1^)- or RIF (0.125 µg mL^-1^)-treated EA.hy926 cells at 24-h intervals between 24 and 120 hpi as described above. The effects of CAM (100–5 µg mL^-1^) and RIF (2–0.125 µg mL^-1^) on host cell fitness was assessed by an MTT assay as described previously (25).

### Plasmid construct

pJHOT::CM*ank13*, which contains *O. tsutsugamushi* codon-optimized chloramphenicol acetyltransferase (CAT) and mScarlet-I genes and a codon-modified *ank13* gene was synthesized by Azenta. The sequence has been deposited to Genbank (accession number PZ464279).

### *O. tsutsugamushi* transformation and outcome assessment

The cassette and flanking regions of pJHOT::CM*ank13* were PCR-amplified using primers Mut3F and Mut3R (Table 1) and Phusion High-Fidelity DNA polymerase (ThermoFisher Scientific) per the manufacturer’s instructions. Thermal cycling conditions consisted of an initial denaturation step of 98°C for 30 s, followed by 34 cycles of 98°C for 10 s, 50°C for 10 s, 72°C for 4 min, and a final extension step at 72°C for 5 min. The resulting amplicons were subjected to agarose gel electrophoresis, after which they were visualized using a Blue View Transilluminator (Vernier Biotechnology) and purified using the NucleoSpin Gel and PCR Clean-up kit (Macherey-Nagel). The purified PCR products were transformed into *O. tsutsugamushi* using either electroporation or CaCl_2_ as follows. One 75-cm^2^ flask containing a confluent monolayer of heavily infected (∼95%) HeLa cells at 3 dpi served as the source of bacteria. Host cell-free organisms were recovered as described above and split into three equal aliquots, two of which were either subjected to electroporation or CaCl_2_-based transformation. The third aliquot served as a no-transformation control. For electroporation transformations, bacteria were washed 3 times with 250 mM sucrose and electroporated with 10 µg of PCR product (3 kV·cm^−1^, 200 Ω, 25 µF, 5.0 ms) as described Kim *et al* (65). For CaCl_2_-based transformation, bacteria were incubated with transformation mix (10 mM Tris, pH 7.4, 50 mM CaCl_2_) and PCR product at room temperature with agitation for 30 min as described by Wan *et al* and Wang *et al* (69, 70). Transformed or non-transformed control *O. tsutsugamushi* were inoculated into a confluent monolayer of EA.hy926 cells in a 25-cm^2^ culture flask. The cells were cultivated at 35°C in a humidified incubator with 5% CO_2_.

To assess the efficiency by which transformed bacteria successfully invaded and survived in host cells until 72 hpi, host cell-free bacteria were recovered from each of the three conditions and DNA isolated as described above. qPCR was performed using primer sets targeting the *O. tsutsugamushi* 16S rRNA gene (*ott16s*) (90), human *GAPDH* (71), *CAT* gene, DT5 transcriptional terminator (Table 1), and the PerfeCTa SYBR Green Fastmix. Thermal cycling conditions were 95°C for 30 s followed by 40 cycles of 95°C for 10 s to 54°C for 10 s, and a 65°C-to-95°C melt curve. Relative expression was determined using the 2^-ΔΔCT^ method (91). Bacterial load was measured as the *ott16s*:*GAPDH* ratio. Transformation efficiency was inferred based on the *cat*:*ott16S* and DT5 *terminator*:*ott16S* ratios. As a complementary approach, DNA was subjected to PCR using the Mut0-F/R and *ott16S*-F/R primer sets. pJHOT::CM*ank13* plasmid and purified *O. tsutsugamushi* DNA served as amplification controls. To compare host cell viability post-incubation with *O. tsutsugamushi* bacteria that had been subjected to electroporation or CaCl_2_-based transformation, phase micrographs were acquired using a TCS SP8 microscope (Leica Microsystems) at 24 hpi.

### Transformant outgrowth and confirmation of allelic exchange

One 75-cm^2^ flask containing a confluent monolayer of heavily infected (∼95%) HeLa cells at 3 dpi served as the source of bacteria. The entire batch of host cell-free *O. tsutsugamushi* bacteria was subjected to CaCl_2_-based transformation and inoculated into a confluent monolayer of EA.hy926 cells in a 75-cm^2^ flask containing complete media without antibiotic. At 72 hpi, CAM was added to a final concentration of 5 µg mL^-1^. The EA.hy926 cells were monitored daily for cytopathic effects. Every 3 d, media was replaced with fresh complete media containing 5 µg mL^-1^ CAM. Every 14 d, media was removed, and 1 ml of 0.5% trypsin (Gibco) was added for 5 min. Two ml of complete media was added for a final volume of 3 ml that was partitioned and analyzed as follows. 0.1 ml was seeded onto a glass coverslip, placed into 24-well plate, and examined 72 h later for infected cells via immunofluorescence microscopy. 1.9 ml was inoculated onto a monolayer of naïve EA.hy926 cells that were ∼50% confluent in a final volume of 10 ml of complete media containing 5 µg mL^-1^ CAM. The remaining 1 ml was homogenized as described above to recover any host cell-free *O. tsutsugamushi* organisms from which DNA was purified using the DNeasy Blood and Tissue kit (Qiagen). To confirm cassette presence and *O. tsutsugamushi* load, PCR was performed using MyTaq polymerase Red (Bioline) and primer sets Mut0-F/R, Mut1-F/R, Mut2-F/R, and *ott16S* F/R primer sets (Table 1). Thermal cycling conditions consisted of an initial denaturing step at 95°C for 1 min, followed by 34 cycles of 95°C for 15 s, 55°C for 15 s, and 72°C for 20 s and a final extension at 72°C for 30 s. Amplicons and the Hyperladder 1kB or 50 bp DNA ladder (Meridian Biosciences) were electrophoresed on an agarose gel and imaged via the ChemiDoc MP Imaging System (BioRad).

To confirm the presence of CM*ank13* and the cassette, PCR was performed using Mut3-F/R (Table 1) and Phusion High-Fidelity DNA polymerase (ThermoFisher Scientific) per the manufacturer’s instructions. Thermal cycling conditions consisted of an initial denaturation step of 98°C for 30 s, followed by 34 cycles of 98°C for 10 s, 50°C for 10 s, 72°C for 5 min, and a final extension step at 72°C for 5 min. To confirm chromosomal insertion of CM*ank13* and the cassette, PCR was performed using Mut4-F/R primer pairs (Table 1) and Platinum SuperFi II DNA polymerase (ThermoFisher Scientific) per the manufacturer’s instructions. Thermal cycling conditions consisted of an initial denaturation step of 98°C for 30 s, followed by 34 cycles of 98°C for 10 s, 60°C for 10 s, 72°C for 2 min, and a final extension step at 72°C for 5 min. Resulting amplicons were subjected to agarose gel electrophoresis, after which they were imaged via the ChemiDoc MP Imaging System, excised, and purified using the NucleoSpin Gel and PCR Clean-up kit. The purified PCR products were sequenced via nanopore amplicon sequencing by Eurofins Genomics. The provided sequences were analyzed using SnapGene software version 7.0.

To detect replacement of wild-type *ank13* with CM*ank13*, PCR was performed using MyTaq polymerase Red (Bioline) and *ank13*-F/R, CM*ank13*-F/R, and *ott16S-*F/R primer sets (Table 1). Thermal cycling conditions consisted of an initial denaturing step at 95°C for 1 min, followed by 34 cycles of 95°C for 15 s, 55°C for 15 s, and 72°C for 10 s and a final extension at 72°C for 10 s. Amplicons and the Hyperladder 1kB or 50 bp DNA ladder (Meridian Biosciences) were electrophoresed on an agarose gel and imaged via the ChemiDoc MP Imaging System (BioRad).

### Statistical analysis

GraphPad Prism version 9 (San Diego, CA) was used for all statistical analyses. One-way analysis of variance ANOVA with Tukey’s or Dunnett’s post hoc test was used to evaluate significant differences among groups. P-values <0.05 were considered statistically significant.

## AUTHOR CONTRIBUTIONS

Paige E. Allen, Conceptualization, Data curation, Formal analysis, Funding acquisition, Investigation, Methodology, Project administration, Visualization, Writing – original draft, Writing – review and editing | Jason R. Hunt, Conceptualization, Data curation, Formal analysis, Investigation, Methodology, Project administration, Visualization, Writing – original draft | Travis J. Chiarelli, Conceptualization, Data curation, Formal analysis, Investigation, Methodology, Visualization, Funding Acquisition, Writing – review and editing | Jason A. Carlyon, Conceptualization, Funding acquisition, Investigation, Methodology, Project administration, Resources, Supervision, Writing – review and editing

## ACKNOWLEDGEMENTS

We are grateful to the many researchers whose pioneering work, much of which is cited herein, set the stage for us to undertake this study. We are also thankful to Dr. Hwan Kim (Renaissance School of Medicine at Stony Brook University) for helpful discussions.

## FUNDING

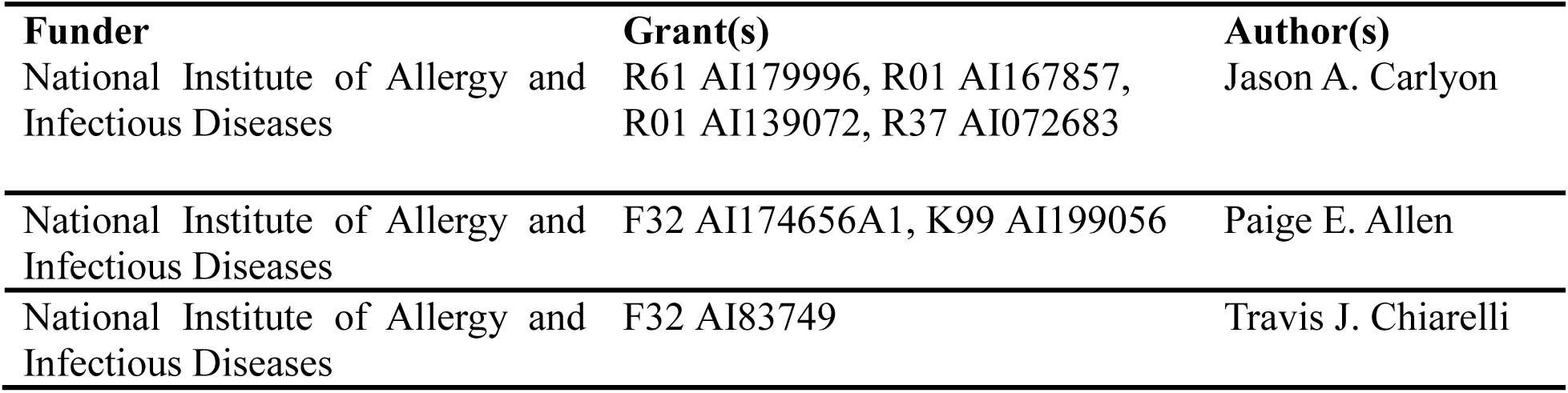

